# Evidence for a potential role of miR-1908-5p and miR-3614-5p in autoimmune disease risk using genome-wide analyses

**DOI:** 10.1101/286260

**Authors:** Inken Wohlers, Lars Bertram, Christina M. Lill

## Abstract

Genome-wide association studies (GWAS) have identified a large number of genetic risk loci for autoimmune diseases. However, the functional variants underlying these disease associations remain largely unknown. There is evidence that microRNA-mediated regulation may play an important role in this context. Therefore, we assessed whether autoimmune disease loci unfold their effects via altering microRNA expression in relevant immune cells.

To this end, we performed microRNA expression quantitative trait loci (eQTL) analyses across 115 GWAS regions associated with 12 autoimmune diseases using next-generation sequencing data of 345 lymphoblastoid cell lines. Statistical analyses included the application and extension of a recently proposed framework (joint likelihood mapping), to microRNA expression data and microRNA target gene enrichment analyses of relevant GWAS data.

Overall, only a minority of autoimmune disease risk loci may exert their pathophysiologic effects by altering miRNA expression based on JLIM. However, detailed functional fine-mapping revealed two independent GWAS regions harboring autoimmune disease risk SNPs with significant effects on microRNA expression. These relate to SNPs associated with Crohn’s disease (CD; rs102275) and rheumatoid arthritis (RA; rs968567), which affect the expression of miR-1908-5p (p_rs102275_=1.44e-20, p_rs968567_=2.54e-14). In addition, an independent CD risk SNP, rs3853824, was found to alter the expression of miR-3614-5p (p=5.70e-7). To support these findings, we demonstrate that GWAS signals for RA and CD were enriched in genes predicted to be targeted by both miRNAs (all with *p*<0.05).

In summary, our study points towards a pathophysiological role of miR-1908-5p and miR-3614-5p in autoimmunity.

## Introduction

The number of risk loci identified by genome-wide association studies (GWAS) has seen a steep increase for many genetically complex diseases over recent years (1). This is especially true for autoimmune diseases where the number of independent GWAS loci has proven to be particularly high (up to more than 100 per disease (www.immunobase.org)). Despite this wealth of newly described genetic susceptibility loci, the precise genes and polymorphisms and, thus, pathophysiological mechanisms underlying these association findings remain largely unknown. Owing to the lack of disease-associated variants with an obvious functional impact, e.g. those changing a gene’s coding sequence, it has been postulated that a sizeable fraction of variants found to be associated by GWAS may unfold their effects by altering gene expression. To test this hypothesis with regard to the expression of messenger RNAs (mRNAs) a recent study (2) developed and applied a novel analytical framework (“joint likelihood mapping” [JLIM]) to GWAS and RNA sequencing data in autoimmune diseases. Despite excellent power, the authors of this study found that probably only about 25% of autoimmune disease GWAS signals act by affecting mRNA levels in cis, i.e. of nearby genes. These data suggest that other, possibly more complex, regulatory mechanisms may underly genetic association findings. To this end, a growing body of functional evidence points towards a central role of microRNAs (miRNAs) in numerous genetically complex human traits and disorders, including autoimmune diseases (for reviews see e.g. ref. (3,4)). MiRNAs are an abundant class of small, non-coding RNAs that post-transcriptionally regulate gene expression in *cis* and *trans* by targeting and degrading mRNA molecules thereby effectively altering mRNA translation and protein expression. Therefore, in this study, we systematically investigated the potential effects of GWAS-derived autoimmune risk variants on miRNA expression measured by next-generation sequencing (RNA-seq) in lymphoblastoid cell lines. One analytical arm of our study entailed the application and extension of the JLIM framework to these miRNA expression data.

## Methods

### Subjects

The RNA-Seq dataset analyzed in this study comprised 345 HapMap (5) individuals of Caucasian descent, for which raw small RNA and messenger RNA sequencing data were available from peripheral lymphoblastoid cell line (LCL) samples generated by the Genetic European Variation in Health and Disease (GEUVADIS) Consortium (6) (Supplementary Tables 1 and 2). Genetic data for these individuals were generated as part of the 1000 Genomes project from whole genome and whole exome sequencing data (see Supplementary Methods for specific data sources).

### Processing of small RNA and mRNA sequencing data

Small RNA sequencing reads were assigned to 2,588 mature human miRNA sequences from miRBase database release 21 using KALLISTO (7). Only miRNAs with estimated read counts (EST) >0 in at least 90% of all samples were included in the subsequent analyses (i.e., 519 miRNAs). MiRNA ESTs were normalized and their variance stabilized using DESEQ2 (8). Messenger RNA sequencing reads were processed using a similar pipeline (Supplementary Methods). We used PEER (9,10) to remove batch effects and hidden covariates from the expression data (Supplementary Methods and Supplementary Figure 1). For eQTL analysis, the resulting miRNA expression values were subjected to inverse normal transformation. See Supplementary Methods for details on annotations, programs and parameter specifications.

### Processing of genotype data

Data on genetic variants were extracted from independently generated whole genome and whole exome sequencing data. Genetic variants with more than two alleles, a minor allele frequency (MAF) less than 0.01 and/or those showing deviation from Hardy-Weinberg equilibrium at α = 1×10^−6^ were excluded resulting in 9,682,130 variants available for subsequent analyses. Principal components (PCs) were calculated using the EIGENSOFT package (11) and used as covariates in the eQTL analyses (Supplementary Methods).

### eQTL analyses in autoimmune diseases

MiRNA cis eQTL analyses were performed for a total of 284 autoimmune risk SNPs located in 115 GWAS regions across 12 autoimmune diseases. This list comprised GWAS index SNPs that showed genome-wide significant association (*p* <5×10^−8^) with autoimmune diseases and that were located within 1 MB distance from the transcription start site of a miRNA. Overall, we assessed the following 12 diseases and corresponding GWAS: ankylosing spondylitis (12), autoimmune thyroid disease (13), celiac disease (14,15), Crohn’s disease (CD) (16), juvenile ideopathic arthritis (17), multiple sclerosis (18,19), narcolepsy (20), primary biliary cirrhosis (21,22), psoriasis (23), rheumatoid arthritis (RA) (24–26), type 1 diabetes (27–29), and ulcerative colitis (30). Summary statistics of corresponding GWAS and ImmunoChip data were downloaded from Immunobase, a GWAS database for autoimmune diseases (http://www.immunobase.org). In addition, we obtained SNPs associated genome-wide significantly with 11 of the 12 autoimmune diseases from the GWAS catalog (1) (https://www.ebi.ac.uk/gwas) and compared them with the SNP list from Immunobase leading to the addition of 134 further SNPs included in subsequent analyses.

MiRNA cis eQTL analyses were performed using matrixEQTL (31) and were based on an additive linear regression model. The first nine (genetic) PCs and sex were used as covariates. The false discovery rate (FDR) was controlled using the Benjamini-Hochberg procedure (Supplementary Methods).

### Statistical assessment of shared genetic effects

We used JLIM (2) for 20 autoimmune disease risk loci that were densely genotyped on the ImmunoChip array. This yielded 40 combinations of a GWAS index SNP and a miRNA located in sufficient proximity to the respective index SNP (i.e., within 1 MB of the lead risk SNP). Permutations for miRNA eQTL association statistics were generated using PLINK2 (https://www.cog-genomics.org/plink2) as described on the JLIM website. For all analyses, JLIM was used applying the same setup and parameter specifications as described in the original study (2) (see their Supplementary Methods).

### MiRNA target gene enrichment analysis

To assess whether any of the differentially expressed miRNAs are potentially involved in disease pathogenesis, we tested for an enrichment of GWAS association signal in their respective target genes. To this end, we first compiled a list of experimentally validated target genes for each miRNA of interest according to miRTarBase(32)) (including those with “strong evidence” and “less strong evidence”). Second, we used the miRWalk2.0 website (33) to compile two lists comprising miRNA target genes that were predicted by i) at least two out of four and ii) by at least three out of four independent miRNA target prediction programs MIRWALK (33), MIRANDA-rel2010 (34), RNA22v2 (35) and TARGETSCAN6.2 (36)), respectively. Enrichment for significant GWAS signals was tested for all three different target gene lists using PASCAL (37).

### Functional fine-mapping

For ***in silico*** fine-mapping of disease loci, we used Ensembl v.78 annotations, regulatory information from Oreganno (38), LDLink (39) which links to RegulomeDB (40), and previously published predicted miRNA promoters (41).

## Results

A total of 115 of the 549 autoimmune disease risk regions (~20%) assessed in this study encompassed (within 1MB) miRNA genes expressed in the lymphoblastoid cell lines, ranging from 13% to 57% per autoimmune GWAS (Supplementary Excel file). Of those, nearly all contained at least one significant miRNA eQTL signal within 1 MB; however, in most instances, the respective index risk SNP and the best miRNA eQTL SNP were not in linkage disequilibrium [LD] Only 14 GWAS index SNPs located in 9 GWAS loci were in at least moderate LD (r^2^>=0.1) with the best miRNA eQTL SNP. Statistical analyses revealed that four GWAS index SNPs in two genome-wide significant autoimmune risk loci, i.e. one region on chromosome 11q.12.2 and one region on chromosome 17q22, showed significant (i.e., FDR at 5%) miRNA eQTL effects (Table 1, and see below). In addition, across the total of four genome-wide significant autoimmune index SNPs, for 36 of 558 miRNA-index SNP combinations, an index SNP had nominally significant effects on miRNA expression, however, these results did not withstand FDR control. In only 10 miRNA-index SNP combinations, a possible effect of a risk variant on miRNA expression was supported by LD of r^2^>=0.1 between index SNP and best miRNA eQTL SNP.

The first locus on chromosome 11q.12.2 (commonly attributed to genes *MYRF*, *FADS1*, *FADS2*) has been implicated in CD (index SNPs rs102275 (16) and rs174537 (42), r^2^ = 0.93) as well as in RA (index SNP rs968567, r^2^ to rs102275 = 0.31) (24). Our analyses indicate that these SNPs are eQTLs for miR-1908-5p (see below). The CD risk allele at rs102275 shows increased miR-1908-5p expression (rs102275: ß = 0.69, *p* = 1.44e-20; Supplementary Figure 2), while the effect of the RA risk allele at rs968567 points in the opposite direction (rs968567: ß = 0.75, *p* = 2.54e-14; Supplementary Figure 3). We observed other eQTL SNPs for miR-1908-5p in this region showing even stronger association with miRNA expression (e.g. rs174544; ß = 0.85, *p* = 5.82e-28). These variants display strong LD with the index CD SNP (r^2^ = 0.79 with rs102275) and weaker LD with the index RA SNP (r^2^ = 0.40), possibly indicating that dysregulated expression of miR-1908-5p may be more prominent in CD than in RA.

In addition to being eQTL loci for miR-1908-5p, the two CD/RA risk SNPs on chromosome 11q.12.2 (i.e. rs102275 and rs968567) are also eQTLs for *FADS1*, *FADS2* and *TMEM258* mRNA expression in the GEUVADIS lymphoblast dataset (6) as well as in a number of other tissues in GTEx (43) (Supplementary Table 3). Interestingly, the genomic region encoding hsa-miR-1908-5p is located in intron1 of the principal isoform of *FADS1* (and exons of other *FADS1* transcripts; Supplementary Figure 4). However, *FADS1* expression is comparatively low in the GEUVADIS dataset (on average nine transcripts per million mapped reads (TPM); Supplementary Figure 5) and alignments of the 5’ and 3’ mature miRNA and *FADS1* sequence reads show that the observed expression of hsa-miR-1908-5p does not represent an artifact due to co-expression of *FADS1* (Supplementary Figure 6). Furthermore, expression analyses of transcription factors predicted to bind in proximity to miR-1908-5p (Supplementary Figure 7) revealed that the expression of *STAT1*, *RB1*, and *IRF1* is significantly negatively correlated with miR-1908-5p expression (Bonferroni corrected p<0.05; Supplementary Figure 8). MiR-1908-5p expression could, thus, possibly be regulated by *STAT1*, *RB1* and/or *IRF1*.

The second locus on chromosome 17q22 (commonly attributed to genes *C17orf67* and *DGKE*) has thus far only been implicated in CD (42). The risk allele at rs3853824 was associated with a modest but highly significant increase of miR-3614-5p expression (ß = - 0.38, *p* = 5.70e-7; Supplementary Figure 9). This variant is in strong LD with the “best” hsa-miR-3614-5p eQTL SNP in this region, i.e. rs35109497 (ß = −0.52, *p* = 1.17e-10; r^2^ to rs3853824 = 0.66). Apart from being a miRNA eQTL, rs3853824 is also associated with mRNA expression of *DGKE* in GEUVADIS and GTEx (2 tissues), as well as of *RP11-670E13.2* in GEUVADIS and of *C17orf67* in GTEx only (2 tissues). MiRNA hsa-miR-3614-5p is moderately expressed in the GEUVADIS dataset (on average 18 TPM; Supplementary Figure 9) and is located in exon 9/9 of the principal isoform of *TRIM25* (Supplementary Figure 10). Analogous to hsa-miR-1908-5p we confirmed that the hsa-miR-3614-5p expression in GEUVADIS does not represent an artifact due to parallel expression of mRNAs in this region (Supplementary Figure 11). *In silico* fine-mapping did not show any transcription factor binding sites close to SNPs of interest according to Oreganno (Supplementary Figure 12). Two polymorphisms (with r^2^ ≥0.8 with rs3853824) were assigned a RegulomeDB score of 2b (Supplementary Figure 13). This indicates transcription factor binding, an identified motif, DNase Footprint and a DNase peak.

If the identified eQTL effects at miR-1908-5p and miR-3614-5p were, indeed, pathogenetically relevant as suggested by our eQTL results, one would expect the targets of one or both miRNAs to be enriched with GWAS SNPs showing association with CD and/or RA. The number of (predicted) target genes for miR-1908-5p and miR-3614-5p varied widely in the three tested settings (i.e. functionally validated targets versus targets predicted by at least two or three methods, respectively; Supplementary Tables 4 and 5). Enrichment analyses showed significant results for all analysis settings for both miRNAs (miR-1908-5p in CD: *p* = 8.98e-9 for at least two predicted targets, *p* = 5.80e-5 for at least three predicted targets, miR-1908-5p in RA: *p* = 4.83e-3 for at least two predicted targets, *p* = 3.81e-3 for at least three predicted targets, miR-3614-5p in CD: *p* = 3.21e-8 for at least two predicted targets, *p* = 1.16e-4 for at least three predicted targets; Supplementary Table 4) except for analyses based on the much smaller lists of functionally validated targets which did not reveal significant enrichments (Supplementary Tables 4 and 5).

Despite these highly consistent findings for miR-1908-5p and miR-3614-5p, overall there is little overlap between investigated GWAS SNPs and miRNA eQTLs. This is supported by joint likelihood computation (by applying JLIM) for 29 of the 298 autoimmune index SNPs across 21 GWAS regions. These SNPs were located near a significant miRNA eQTL signal and the regions were sufficiently densely genotyped on the Immunochip array and could thus be formally statistically assessed for a shared effect on autoimmune disease risk and miRNA expression. In accordance with the original study (2), we observe a bimodal distribution of JLIM p-values (Supplementary Figure 14). None of the 40 assessed miRNA-index SNP combinations was significant after multiple testing correction. Only one was nominally significant, but did not survive correction for multiple testing (Supplementary Table 6). Importantly, the two regions on chromosomes 11 and 17 showing significant miRNA eQTL effects for the index risk SNPs were not densely genotyped on the Immunochip array and thus could not be assessed using JLIM.

## Discussion

In the current study, we systematically analyzed all currently established GWAS-derived autoimmune risk SNPs for their potential impact on miRNA expression. While in general the causal contribution of dysregulated miRNA expression in the studied autoimmune diseases appears to be modest as suggested by the JLIM-based analyses, in line with similar analyses performed earlier for mRNA expression patterns, we identified significant eQTL effects for two risk loci on chromosomes 11q.12.2 and 17q22. Overall, our results suggest that the corresponding miRNAs, miR-1908-5p and miR-3614-5p, may play a significant role in CD and/or RA pathogenesis

One of the two miRNAs highlighted in our study, hsa-miR-1908-5p, was recently selected for a blood-based disease classification panel from a total of 863 miRNAs as one of sixteen miRNAs that help to distinguish CD patients from healthy controls and ulcerative colitis patients using a machine learning algorithm (44). This supports our findings that hsa-miR-1908-5p expression is indeed altered in CD. In ancillary analyses, we identified a negative correlation between several immunologically relevant transcription factors (such as STAT1 and RB1) and hsa-miR-1908-5p (see results). Notably, for the RA index SNP rs968567, Lattka et al. predicted binding for STAT1 in the presence of the protective T allele (45). This is in agreement with the miRNA eQTL effect observed here for this SNP in the GEUVADIS data. Additional regulatory effects may be exerted by transcription factor RB1 for which an Oreganno regulatory region is located only 3 bp upstream of rs968567. Furthermore, rs968567 is located within 148 bp of a possible miRNA promoter predicted by Marsico et al.(41). Collectively and consistently, these data point to a regulatory effect of RA risk SNP rs968567 on hsa-miR-1908-5p expression. The potential mechanisms behind the observed eQTL effects of SNPs in this region associated with CD are less obvious as the CD SNP rs102275 displays only modest LD with the RA SNP rs968567 and does not map into the vicinity of regulatory regions containing STAT1, IRF1 or RB1 transcription factor binding sites. A STAT1 targeted regulatory region covering mir-1908 pre-miRNA structure and harboring SNP rs174561, a SNP in LD of r2=1 with the best eQTL SNP rs174544 (r^2^=0.79 with CD index SNP rs102275) may play a role. A similar observation was made for our second CD locus on chromosome 17 near miR-3614-5p for which we delineate converging evidence of an involvement in CD by GWAS and eQTL, but currently lack any clear mechanistic data such as altered binding of a specific transcription factor possibly leading to the observed expressional differences identified in the eQTL analyses.

Despite the compelling agreement of results across multiple domains suggesting the presence of allele-specific expressional changes of hsa-miR-1908-5p and miR-3614-5p in CD and/or RA, it needs to be emphasized that based on the analyses performed in this study we cannot draw any firm conclusions that these effects are, indeed, pathogenetically relevant. For both loci, the autoimmune risk SNPs also represent eQTLs for other genes in that region (see results). Thus, the pathophysiological mechanisms underlying the genetic associations at these two loci could be due to the differential risk-allele specific expression of one or more of the protein-coding genes, the differential expression of the miRNAs as described in this study, or both. Additional limitations of our study relate to the following points: First, the GWAS results included here originate from different primary studies with differing statistical power (depending on sample size) and differing genetic resolution (depending on the microarray employed). Thus, if there is association data for a larger number of variants within any given locus, it is more likely that a miRNA eQTL in the region is also directly assessed within the GWAS. However, this problem is mitigated by the fact that we were able to calculate the correlation (via LD) between variants regardless of whether or not they were directly tested in the primary GWAS. Second, our miRNA eQTL analyses are based on only 345 individuals and only variants with 1% minor allele frequency were included. Increasing the number of individuals would not only increase the statistical power of our eQTL analyses but also allow to assess variants with frequencies <1%. Since we here used previously generated DNA- and RNA-seq data for the eQTL arm of our study, i.e. that of the GEUVADIS consortium, the sample size for these analyses was “fixed”. Only future studies in larger samples will be able to effectively address this limitation. Third, for the eQTL analyses we utilized RNA-seq data generated in lymphoblast cell lines. While this cell type represents a reasonable “surrogate tissue” to study miRNA effects in autoimmune diseases, expressional changes in other pathophysiologically relevant immune cells could not be analyzed here. These can only be detected in efforts generating miRNA (and mRNA) expression profiles in additional cell types allowing a more differentiated and complete overview of the potential transcriptional (dys)regulation in autoimmune diseases.

In conclusion, in this study we systematically assessed whether and which autoimmune disease-associated GWAS variants influence miRNA expression in lymphoblast cell lines. While, overall, the impact of GWAS SNPs on the expressional regulation of miRNAs appears modest, we identified two miRNAs that likely play a role in RA and CD. One of the highlighted miRNAs, i.e. hsa-miR-1908-5p, had already been previously identified to be useful for a molecular diagnosis of CD, lending further support to the conclusions reached by our analyses. Future studies generating miRNA and mRNA transcriptomics data in other cell types are needed to provide a more complete insight into the potential impact of these and other GWAS signals on gene expression relevant in autoimmune diseases.

## Acknowledgements

We thank Lukas Duchrow for support in implementing the enrichment analysis and Colin Schulz for support in paper formatting. We thank Dr. Sung Chun and Prof. Chris Cotsapas for help in applying the JLIM software and Dr. Ashley Beecham for providing additional information regarding the Immunochip array. Computational support was provided by the OMICS compute cluster at the University of Lübeck. I.W. was supported by the Peter und Traudl Engelhorn Foundation. C.M.L. received funding from the German Research Foundation (DFG; FOR2488/1, GZ LI 2654/2-1), the Possehl Foundation, the Renate Maaß Foundation, and the University of Luebeck (section of medicine, J21-2016).

