## Supplementary Materials for "Evidence for a potential role of miR-1908-5p and miR-3614-5p in autoimmune disease risk using genome-wide analyses" (from another version of this preprint: version 1, 2018-03-22)

### Supplementary Methods

**RNA-Seq and genotype data sources**: GEUVADIS raw data files are accessible via the European Nucleotide Archive (http://www.ebi.ac.uk/ena) under accession numbers "ERP001942" and "ERP001941". Read counts and eQTLs of the original study are available via the ArrayExpress database [https://www.ebi.ac.uk/arrayexpress/; accession numbers E-GEUV-1 and E-GEUV-2]. The most recent whole-genome sequencing-based genotype data generated by the 1000 Genomes (1000G) project (GRCh38; phase 3, release date: 20130502) in VCF format was obtained at ftp://ftp.1000genomes.ebi.ac.uk/vol1/ftp/release/20130502/supporting/GRCh38_positions/.

**Computation of principal components**: Processing of the 1000G data was performed using VCFTOOLS (1). In order to calculate principal components (PC), insertions and deletions and variants within regions of high linkage disequilibrium (LD) (2) were removed from the dataset and LD pruning (--indep-pairwise 1000 10 0.05) was performed using PLINK v.1.07 (3). PCs were calculated using SMARTPCA v.13050 from the Eigensoft package (4) with default parameters. Subsequently, Tracy-Widom statistics for determining (unsupervised) influential PCs (4) were computed, and, as a result, the first nine PCs were used as covariates in all subsequent analyses.

**Processing of small RNA and mRNA sequencing data**: MiRNA sequences were downloaded for miRBase database release 21 from ftp://mirbase.org/pub/mirbase/21/mature.fa.gz. For mapping, KALLISTO version 0.42.4 (5) with the options --single, --fragment-length=150, -s 15, number of bootstraps set to 100. Default settings were used for all other parameters. As expected, we observed a high correlation between the miRNA read counts that we estimated from the raw data and the miRNA counts made publicly available in the original study (6) (r=0.99 across all miRNAs investigated in both studies). Reads of mRNAs were mapped to 207,167 coding and non-coding transcript sequences (obtained from the Ensembl database v.78; ftp://ftp.ensembl.org/pub/release-78/fasta/homo_sapiens/{cdna,ncrna}) using KALLISTO (v. 0.42.4 (5)) with default parameters and with the number of bootstraps set to 100. Between-sample normalizations of estimated read counts (EST) and of transcripts per million mapped reads (TPM) were performed using DESEQ2 (7). For batch effect removal, the program PEER (8,9) was used with 10 factors for miRNAs and 20 factors for transcripts. PC analysis showed that for both miRNA and transcripts, expression clustering by either laboratory or population was barely noticeable after removing hidden covariates from the dataset (Supplementary Figure 1).

**Comparison of our miRNA eQTLs to GEUVADIS miRNA eQTLs**: The *ß* of our eQTL linear regression analyses showed a high correlation with the r-value of the publicly available GEUVADIS eQTLs (6) (Pearson's r=0.95). In addition, the effect directions for all miRNA eQTLs obtained in our and in the GEUVADIS analysis matched. Thus, our analysis results are comparable to the GEUVADIS results, with the following differences: We (i) use updated miRNA annotations and SNP IDs, (ii) are more stringent on the number of samples in which a miRNA needs to be expressed to be included in eQTL analysis, and (iii) use genotypes obtained from whole-genome sequencing instead of imputed genotypes for some individuals.

### Supplementary Tables

| population | samples |
|---|---|
| CEU | 84 |
| FIN | 90 |
| GBR | 84 |
| TSI | 87 |

**Table 1: Strata of European ethnicity included in the GEUVADIS dataset.**

| laboratory | mRNA samples | miRNA samples |
|---|---|---|
| CNAG_CRG | 73 | 70 |
| HMGU | 34 | 37 |
| ICMB | 45 | 45 |
| LUMC | 31 | 32 |
| MPIMG | 47 | 47 |
| UNIGE | 85 | 83 |
| UU | 30 | 31 |

**Table 2: Sequencing centers contributing GEUVADIS mRNA and miRNA samples.**

| | FADS1 | FADS2 | TMEM285 | FADS3 | RP11-467L20.10 | BEST1 | DAGLA |
|---|---|---|---|---|---|---|---|
| rs968567 | $2^{*}$ | $22^{*+}$ | $2^{*}$ | 0 | 0 | 0 | 0 |
| rs174537 | $10^{*}$ | $16^{*+}$ | $6^{*}$ | 1 | 1 | 1 | 0 |
| rs102275 | $8^{*}$ | $12^{*+}$ | $3^{*}$ | 1 | 1 | 0 | 0 |
| rs174561 | $12^{*}$ | $16^{*+}$ | $6^{*}$ | 1 | 1 | 0 | 1 |

**Table 3: The number of tissues for which the CD and RA index SNPs are eQTLs for a given gene within analysis of the GTEx data.** The CD and RA index SNPs also represent miRNA eQTLs for hsa-miR-1908-5p. An asterisk $^{*}$ denotes that there is at least one exon eQTL for this gene in the analysis of the GEUVADIS data, a plus sign $^{+}$ denotes gene eQTLs in the analysis of the GEUVADIS data.

| | | | CD | | RA | |
|---|---|---|---|---|---|---|
| methods | conc. | targets | p $X^2$ | p emp | p $X^2$ | p emp |
| miRTarBase | - | 95 | 6.58e-2 | 7.38e-2 | 1.56e-1 | 3.67e-1 |
| Mirwalk/Miranda/rna22/Targetscan | 2 | 3734 | **8.98e-9** | **4.19e-5** | **4.83e-3** | **2.77e-3** |
| Mirwalk/Miranda/rna22/Targetscan | 3 | 1333 | **5.80e-5** | **2.72e-3** | **3.81e-3** | **1.98e-2** |

**Table 4: PASCAL $X^2$ and empirical p-values for the enrichment of GWAS associations for Crohn's disease (CD) and rheumatoid arthritis (RA) in miR-1908-5p target genes.** Significant p-values (α=0.05) are highlighted in bold. Term "conc." denotes the number of methods that need to concordantly predict a target gene in order to include it in the enrichment analysis. Term ”targets” refers to the number of target genes obtained by the corresponding method(s).

| methods | conc. | targets | p $X^2$ | p emp |
|---|---|---|---|---|
| miRTarBase | - | 167 | 5.03e-1 | 5.68e-1 |
| Mirwalk/Miranda/rna22/Targetscan | 2 | 5417 | **3.21e-8** | **1.19e-5** |
| Mirwalk/Miranda/rna22/Targetscan | 3 | 1982 | **1.16e-4** | **2.36e-3** |

**Table 5: PASCAL $X^2$ and empirical p-values for the enrichment of GWAS associations for Crohn's disease in miR-3614-5p target genes.** See Table 4 for legend.

| GWAS lead SNP | | | MiRNA | | eQTL SNP | | | LD | JLIM |
|---|---|---|---|---|---|---|---|---|---|
| SNP | Pos | P | MiRNA | Dist lead | SNP | Pos | P | LD [R2] | P JLIM |
| **Type 1 diabetes** | | | | | | | | | |
| *Onengut-Gumuscu et al.* (10) | | | | | | | | | |
| rs56994090 | 14:100840110 | 1.13e-11 | hsa-miR-345-5p | 532,234 | rs79160867 | 14:100837091 | 7.38e-03 | 0.04 | 0.84 |
| rs402072 | 19:46715865 | 4.72e-08 | hsa-miR-769-5p | 696,904 | rs11668878 | 19:46765116 | 3.08e-03 | 0.03 | 0.81 |
| rs679574 | 19:48702851 | 5.22e-14 | hsa-miR-150-5p | 797,981 | rs11880281 | 19:48671478 | 5.00e-03 | 0.15 | 0.75 |
| rs34536443 | 19:10352442 | 4.39e-15 | hsa-miR-199a-3p | 464,987 | rs91755 | 19:10362894 | 2.71e-03 | 0.06 | 0.77 |
| rs4820830 | 22:30135102 | 1.20e-12 | hsa-miR-3653-3p | 801,926 | rs5997592 | 22:30213107 | 8.14e-04 | 0.09 | 0.98 |
| rs4820830 | 22:30135102 | 1.20e-12 | hsa-miR-3653-5p | 801,876 | rs79955051 | 22:30046891 | 5.15e-03 | 0.03 | 0.79 |
| rs229533 | 22:37191071 | 1.79e-08 | hsa-miR-659-5p | 656,661 | rs71317032 | 22:37199116 | 3.47e-03 | 0.27 | 0.83 |
| rs6476839 | 9:4290823 | 1.03e-09 | hsa-miR-101-3p | 559,522 | rs117421312 | 9:4261744 | 9.62e-04 | 0.03 | 0.95 |
| **Rheumatoid arthritis** | | | | | | | | | |
| *Eyre et al.* (11) | | | | | | | | | |
| rs34536443 | 19:10352442 | 2.70e-13 | hsa-miR-199a-3p | 464,987 | rs91755 | 19:10362894 | 2.71e-03 | 0.06 | 0.78 |
| **Celiac disease** | | | | | | | | | |
| *Trynka et al.* (12) | | | | | | | | | |
| rs4821124 | 22:21625000 | 5.72e-11 | hsa-miR-130b-3p | 28,354 | rs447001 | 22:21656629 | 8.69e-05 | 0.13 | 1 |
| rs4821124 | 22:21625000 | 5.72e-11 | hsa-miR-130b-5p | 28,316 | rs447001 | 22:21656629 | 1.59e-03 | 0.13 | 1 |
| rs4821124 | 22:21625000 | 5.72e-11 | hsa-miR-301b-3p | 28,025 | rs384262 | 22:21658483 | 3.91e-03 | 0.04 | 0.96 |
| rs76830965 | 3:159919889 | 2.57e-27 | hsa-miR-15b-5p | 484,718 | rs33052 | 3:159882300 | 2.94e-03 | 0.05 | 0.8 |
| rs76830965 | 3:159919889 | 2.57e-27 | hsa-miR-16-2-3p | 484,908 | rs2243130 | 3:159993205 | 1.99e-03 | 0.00 | 0.75 |
| rs76830965 | 3:159919889 | 2.57e-27 | hsa-miR-16-5p | 484,865 | rs79300081 | 3:159910520 | 2.71e-03 | 0.01 | 0.86 |
| **Multiple sclerosis** | | | | | | | | | |
| *Beecham et al.* (13) | | | | | | | | | |
| rs6677309 | 1:116537544 | 1.45e-28 | hsa-miR-320b | 134,243 | rs116610628 | 1:116577119 | 2.21e-03 | 0.01 | 0.91 |
| rs34383631 | 11:61025858 | 5.69e-10 | hsa-miR-1908-5p | 789,351 | rs72928583 | 11:61046564 | 8.26e-03 | 0.09 | 0.59 |
| rs1800693 | 12:6330843 | 6.92e-16 | hsa-miR-200c-3p | 632,899 | rs113559360 | 12:6312886 | 2.76e-03 | 0.01 | 0.86 |
| rs4796791 | 17:42378745 | 1.81e-08 | hsa-miR-5010-3p | 135,522 | rs113366589 | 17:42364598 | 7.13e-03 | 0.05 | 0.74 |
| rs1014486 | 3:159973324 | 1.16e-09 | hsa-miR-15b-5p | 431,283 | rs33052 | 3:159882300 | 2.94e-03 | 0.00 | 0.79 |
| rs1014486 | 3:159973324 | 1.16e-09 | hsa-miR-16-2-3p | 431,473 | rs2243130 | 3:159993205 | 1.99e-03 | 0.02 | 0.74 |
| rs1014486 | 3:159973324 | 1.16e-09 | hsa-miR-16-5p | 431,430 | rs79300081 | 3:159910520 | 2.71e-03 | 0.04 | 0.56 |
| **Primary biliary cirrhosis** | | | | | | | | | |
| *Liu et al.* (14) | | | | | | | | | |
| rs1800693 | 12:6330843 | 1.18e-14 | hsa-miR-141-3p | 633,312 | rs11064135 | 12:6305745 | 2.25e-03 | 0.07 | 0.85 |
| rs1800693 | 12:6330843 | 1.18e-14 | hsa-miR-200c-3p | 632,899 | rs11831297 | 12:6381443 | 4.57e-03 | 0.00 | 0.59 |
| rs11117433 | 16:85985910 | 1.41e-09 | hsa-miR-1910-5p | 244,241 | rs72807051 | 16:85972327 | 2.11e-03 | 0.07 | 0.92 |
| rs11117433 | 16:85985910 | 1.41e-09 | hsa-miR-6774-3p | 67,514 | rs9674233 | 16:85978993 | 1.60e-03 | 0.07 | 0.95 |
| rs34536443 | 19:10352442 | 1.23e-12 | hsa-miR-199a-3p | 464,987 | rs91755 | 19:10362894 | 2.71e-03 | 0.06 | 0.79 |
| rs522127 | 3:160016881 | 2.98e-22 | hsa-miR-16-2-3p | 387,916 | rs2243130 | 3:159993205 | 1.99e-03 | 0.00 | 0.91 |
| rs35188261 | 7:129043485 | 6.52e-22 | hsa-miR-182-5p | 726,962 | rs4728152 | 7:129098490 | 7.17e-04 | 0.03 | 0.99 |
| rs35188261 | 7:129043485 | 6.52e-22 | hsa-miR-183-5p | 731,482 | rs1077482 | 7:129103017 | 3.96e-05 | 0.04 | 1 |
| **Ankylosing spondylitis** | | | | | | | | | |
| *Cortes et al.* (15) | | | | | | | | | |
| rs1860545 | 12:6337611 | 2.78e-10 | hsa-miR-200c-3p | 626,131 | rs113559360 | 12:6312886 | 2.76e-03 | 0.01 | 0.91 |
| rs35164067 | 19:10414505 | 3.43e-10 | hsa-miR-199a-3p | 402,924 | rs13346189 | 19:10505968 | 4.51e-04 | 0.01 | 0.9 |
| rs1128905 | 9:136359387 | 6.95e-09 | hsa-miR-126-5p | 311,229 | rs117893486 | 9:136419135 | 7.87e-03 | 0.05 | 0.64 |
| **Narcolepsy** | | | | | | | | | |
| *Faraco et al.* (16) | | | | | | | | | |
| rs10995245 | 10:62631615 | 4.81e-08 | hsa-miR-1296-5p | 741,397 | rs113879650 | 10:62689017 | 4.56e-03 | 0.16 | 0.17864 |
| **Psoriasis** | | | | | | | | | |
| *Tsoi et al.* (17) | | | | | | | | | |

| | | | | | | | | | |
|---|---|---|---|---|---|---|---|---|---|
| rs117135073 | 10:62595406 | 2.20e-12 | hsa-miR-1296-5p | 777,606 | rs113879650 | 10:62689017 | 4.56e-03 | 0.00 | 0.75 |
| rs77520588 | 1:116744732 | 5.94e-15 | hsa-miR-320b | 72,945 | rs77520588 | 1:116744732 | 2.76e-03 | 1.00 | 0.01167 |
| rs10744705 | 12:6389529 | 1.69e-08 | hsa-miR-200c-3p | 574,213 | rs11831297 | 12:6381443 | 4.57e-03 | 0.00 | 0.88 |
| rs34536443 | 19:10352442 | 9.01e-14 | hsa-miR-199a-3p | 464,987 | rs91755 | 19:10362894 | 2.71e-03 | 0.06 | 0.77 |
| rs3135952 | 3:48468811 | 2.46e-17 | hsa-miR-191-3p | 551,820 | rs73074378 | 3:48404830 | 7.25e-03 | 0.01 | 0.53 |
| rs3135952 | 3:48468811 | 2.46e-17 | hsa-miR-425-5p | 551,388 | rs4858795 | 3:48473670 | 1.68e-03 | 0.04 | 0.11992 |

**Table 6: Loci and miRNAs assessed with JLIM to find statistical evidence for a shared genetic effect.** Only immunochip-based GWAS data from loci that are densely genotyped on the immunochip were used.

### Supplementary Figures

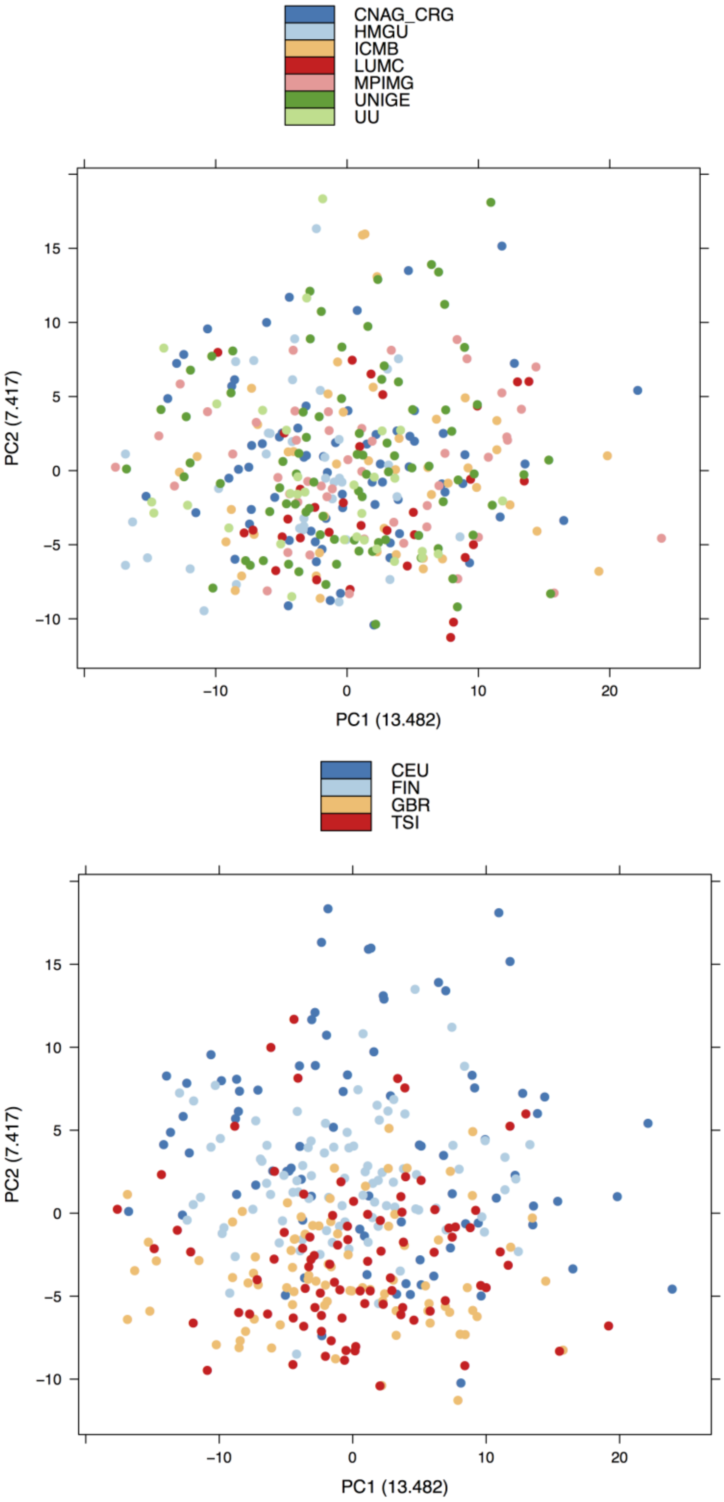

**Figure 1: PC analysis of between-sample normalized, DESEQ2 variance-stabilized miRNA counts**. Samples are colored according to lab and population.

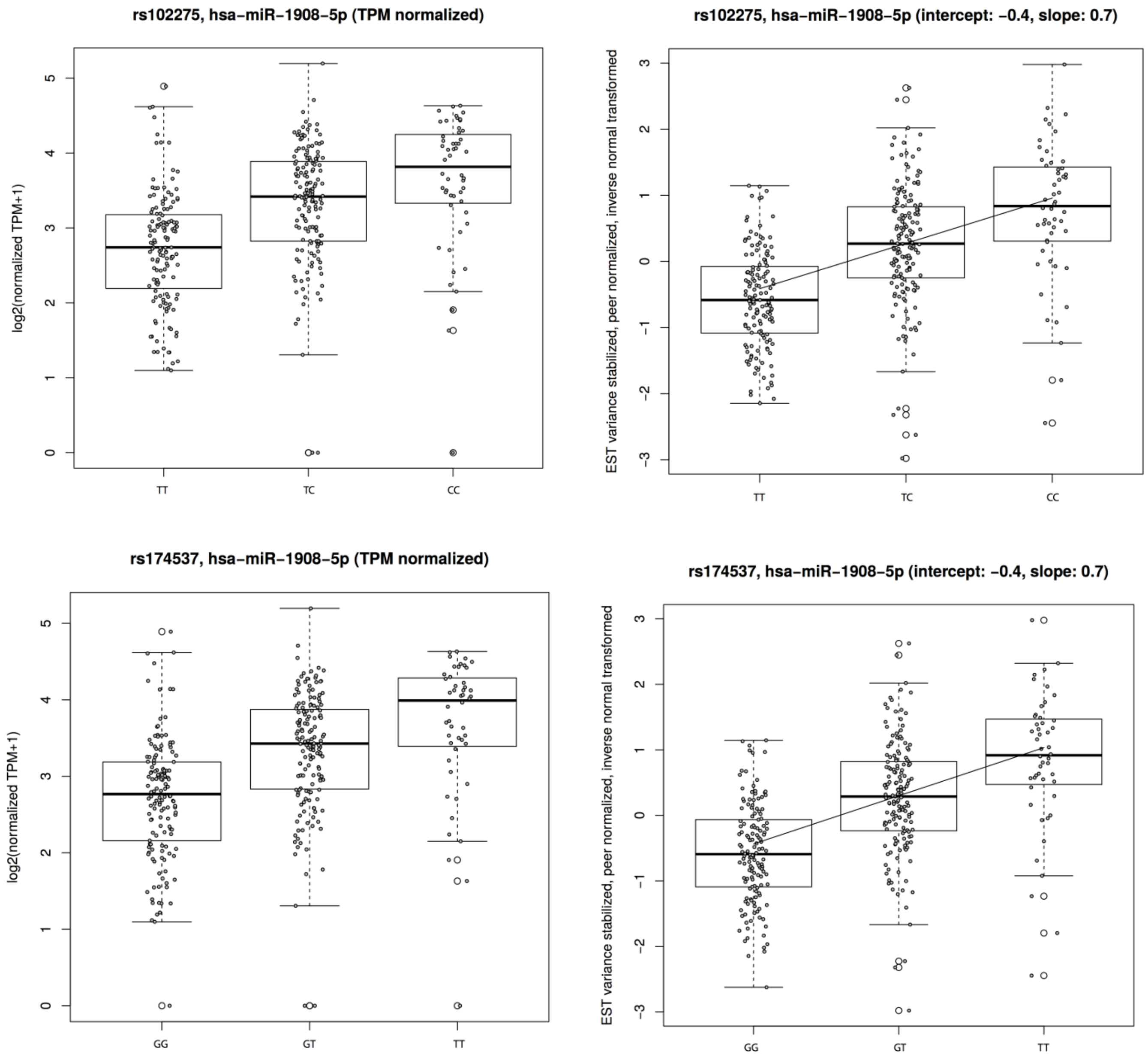

**Figure 2: hsa-miR-1908-5p gene expression levels according to genotypes of Crohn's disease risk SNPs rs102275 and rs174537.** Left: Boxplots of expression data quantified in transcripts per million (TPM), stratified by genotype. Right: Boxplots of processed expression used for eQTL analysis.

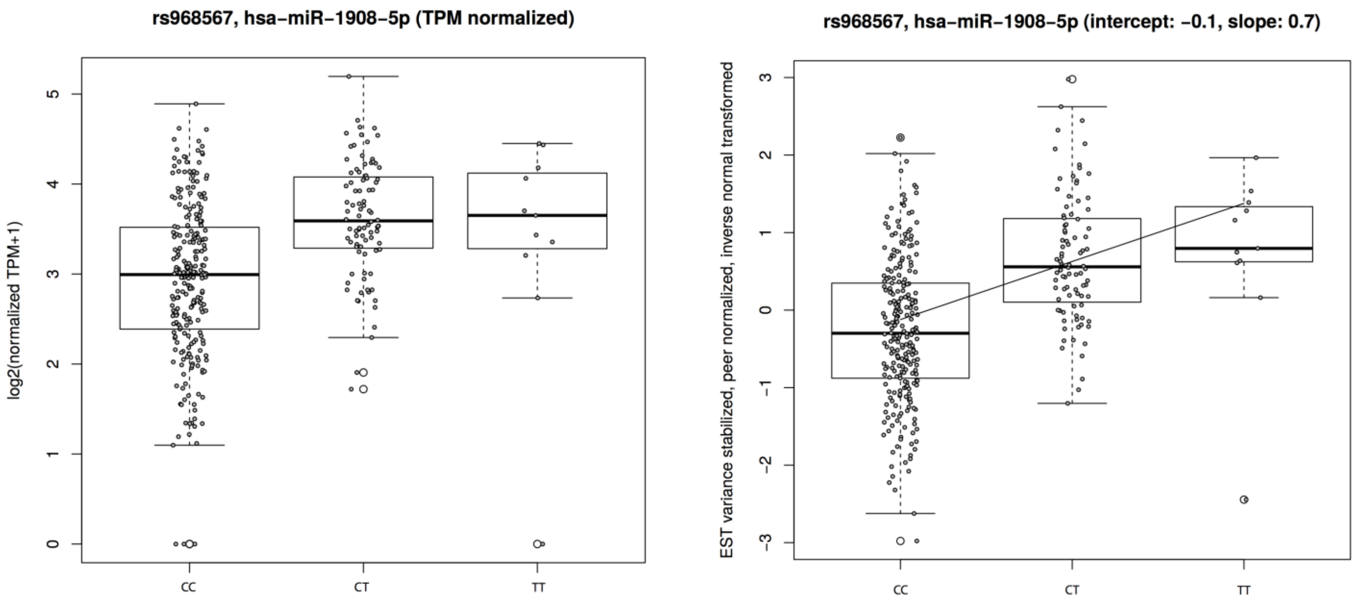

**Figure 3: hsa-miR-1908-5p gene expression levels according to genotypes of rheumatoid arthritis risk SNP rs968567.** See Suppl. Figure 2 for legend.

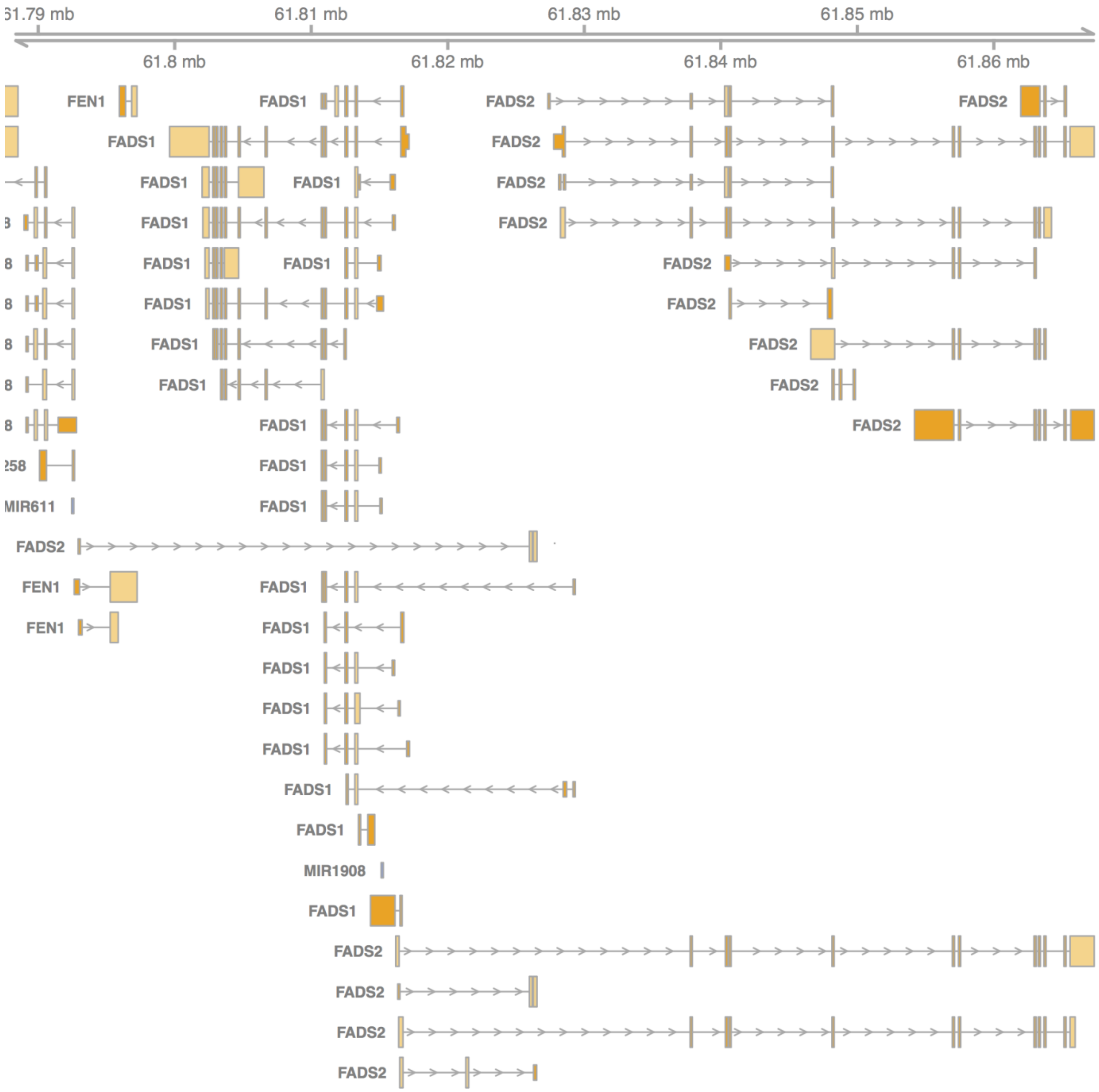

**Figure 4: Region of transcripts that overlap mir-1908.** Narrow boxes denote untranslated regions.

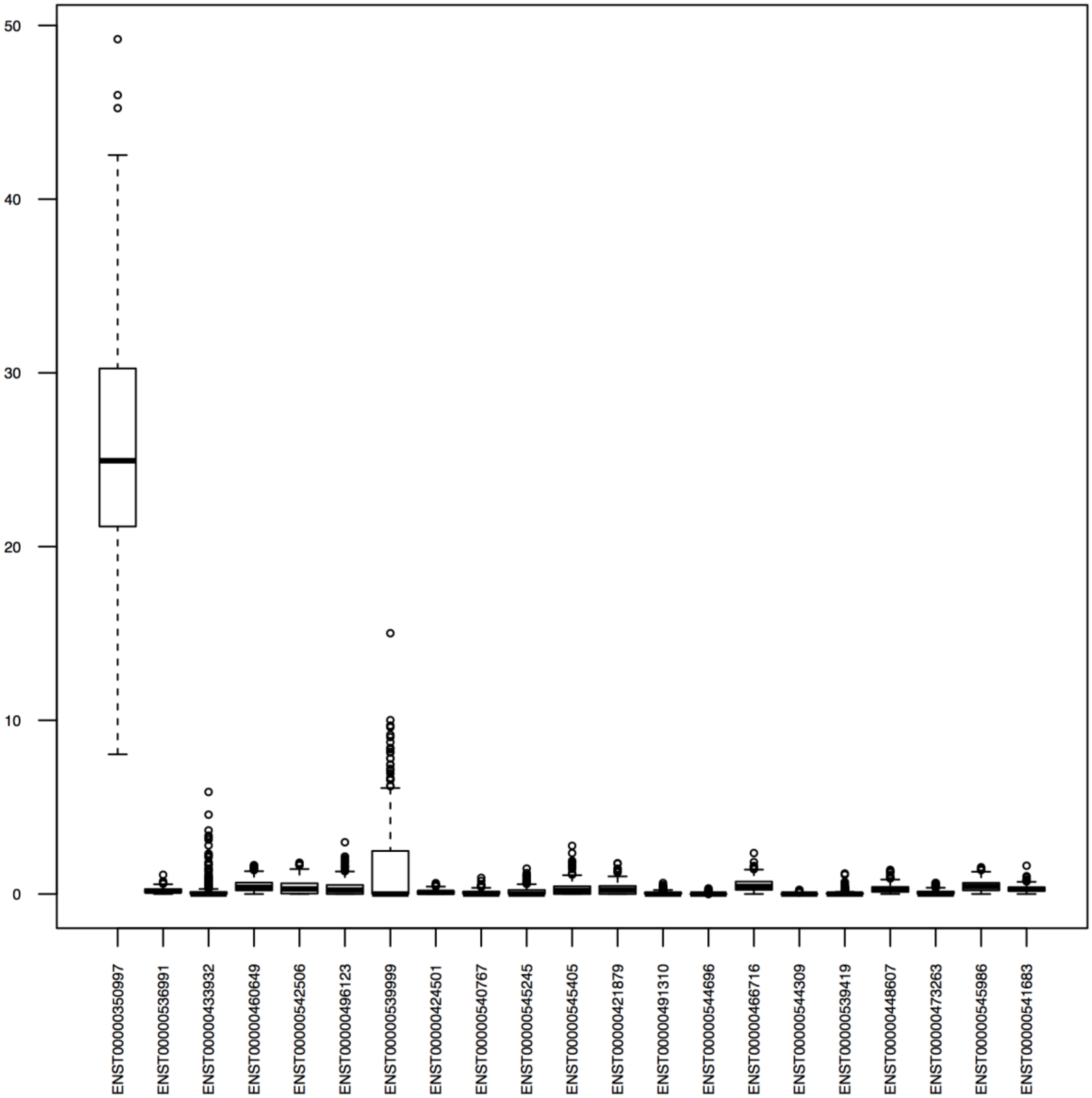

**Figure 5: Expression of FADS1 isoforms in transcripts per million mapped reads (TPM).**

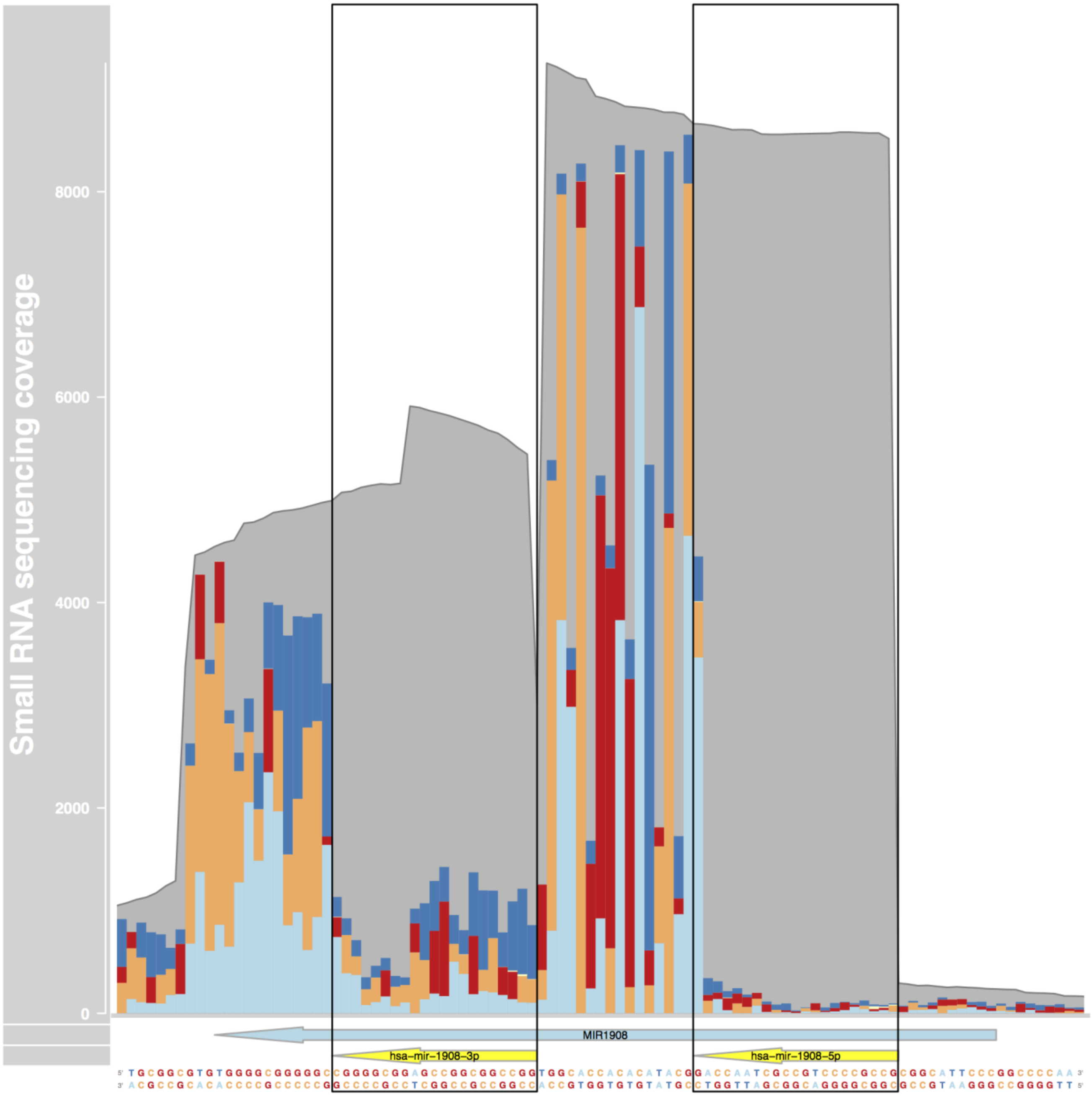

**Figure 6: Small RNA sequencing reads mapped to mir-1908 including the mature hsa-miR-1908-5p (right) and hsa-miR-1908-3p (left) miRNAs and to its surrounding sequence.** The mismatches downstream of the mature miRNA sequences can be attributed to Illumina primers and adapters (as the read length is 36 exceeding miRNA length).

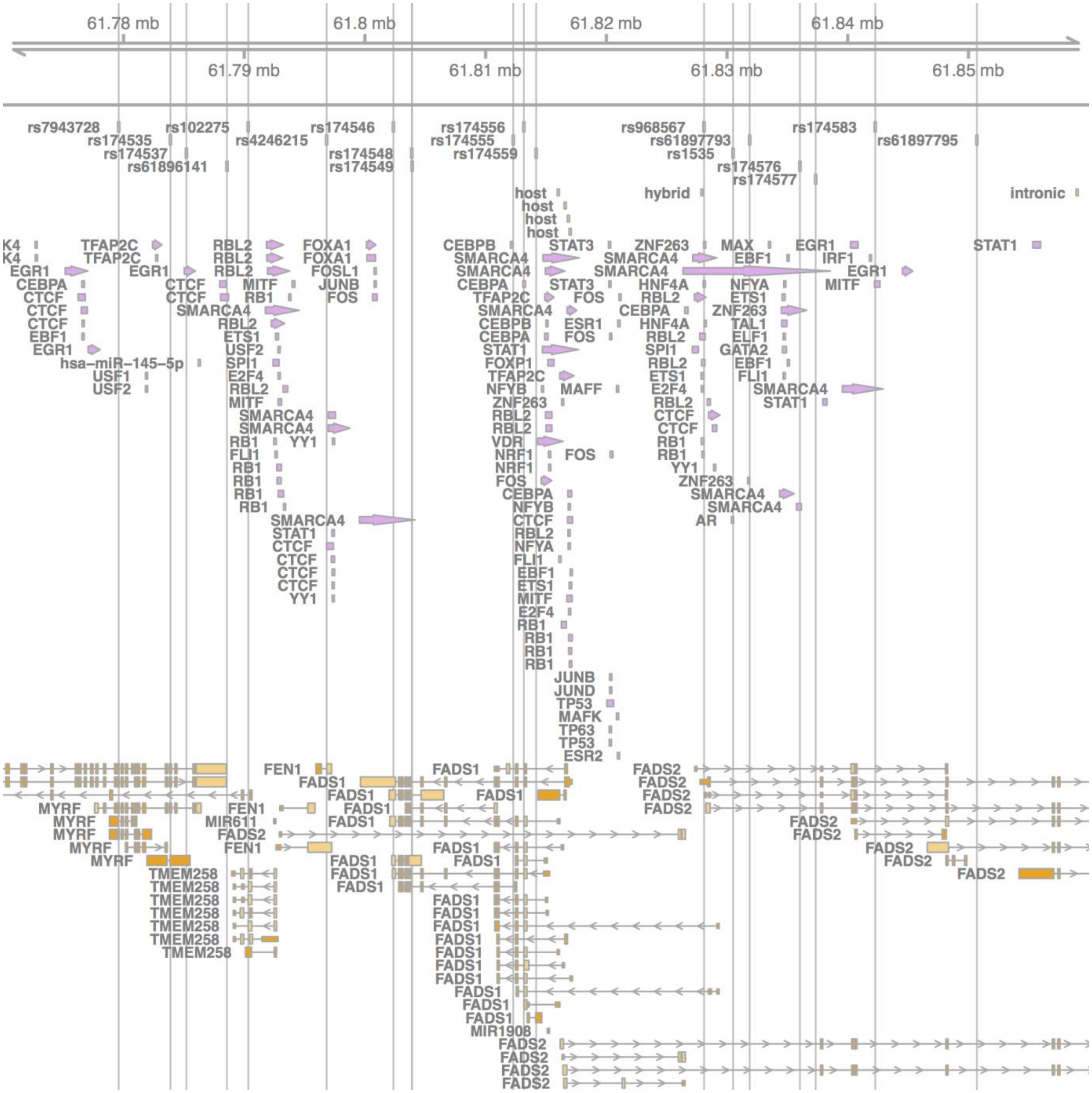

**Figure 7: Visualization of GWAS SNPs that are miRNA eQTLs together with overlapping transcripts.** Regulatory regions from Oreganno are labeled with corresponding targeting transcription factors (pink) and miRNA promoters predicted by Marsico et al. (18)

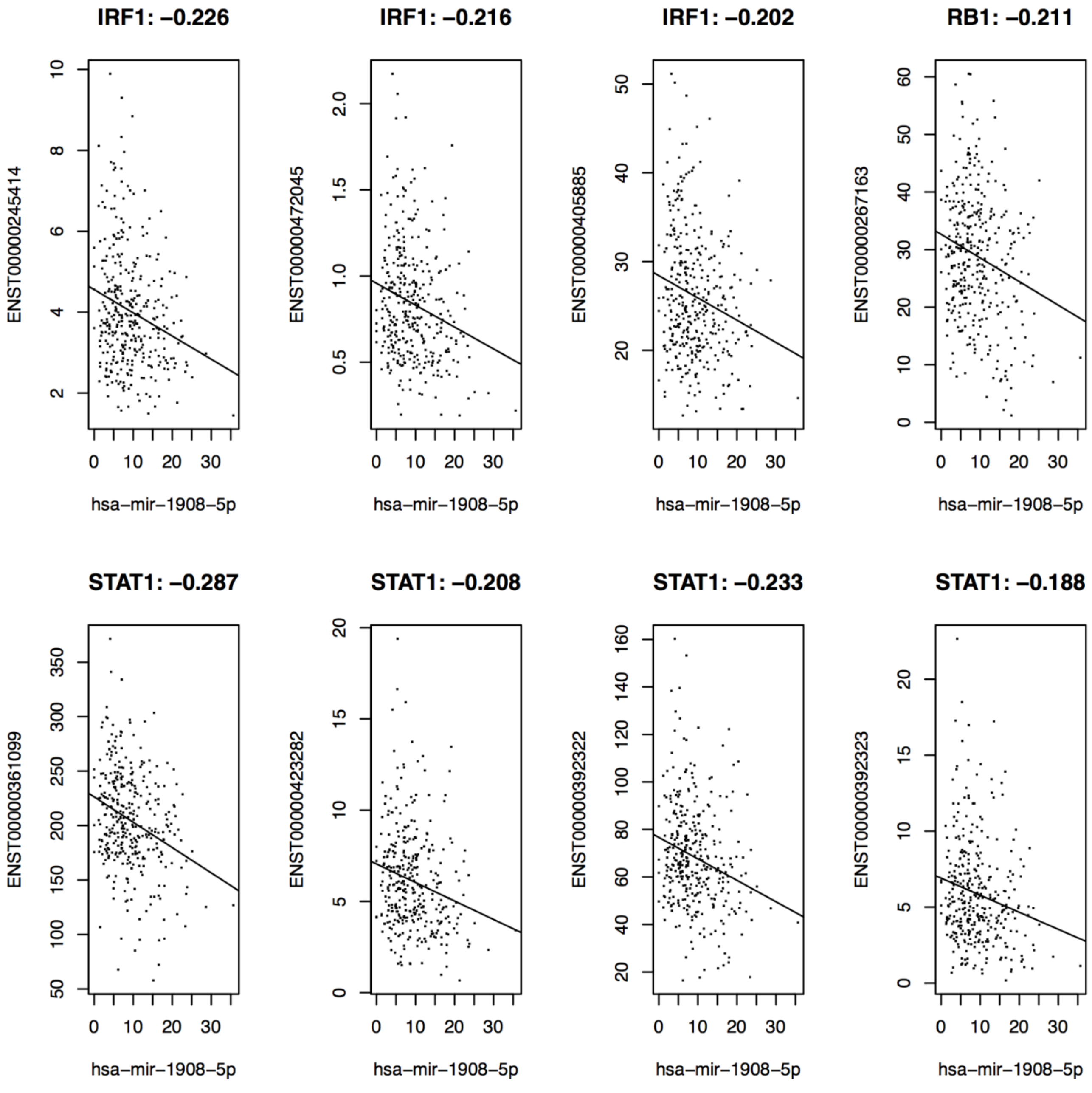

**Figure 8: Expression of transcription factors (in TPM) predicted to target the region surrounding SNP rs968567 correlated with the corresponding miRNA expression.** The correlation is reported in the title of each figure, only significant transcripts with p-value <0.01 are depicted.

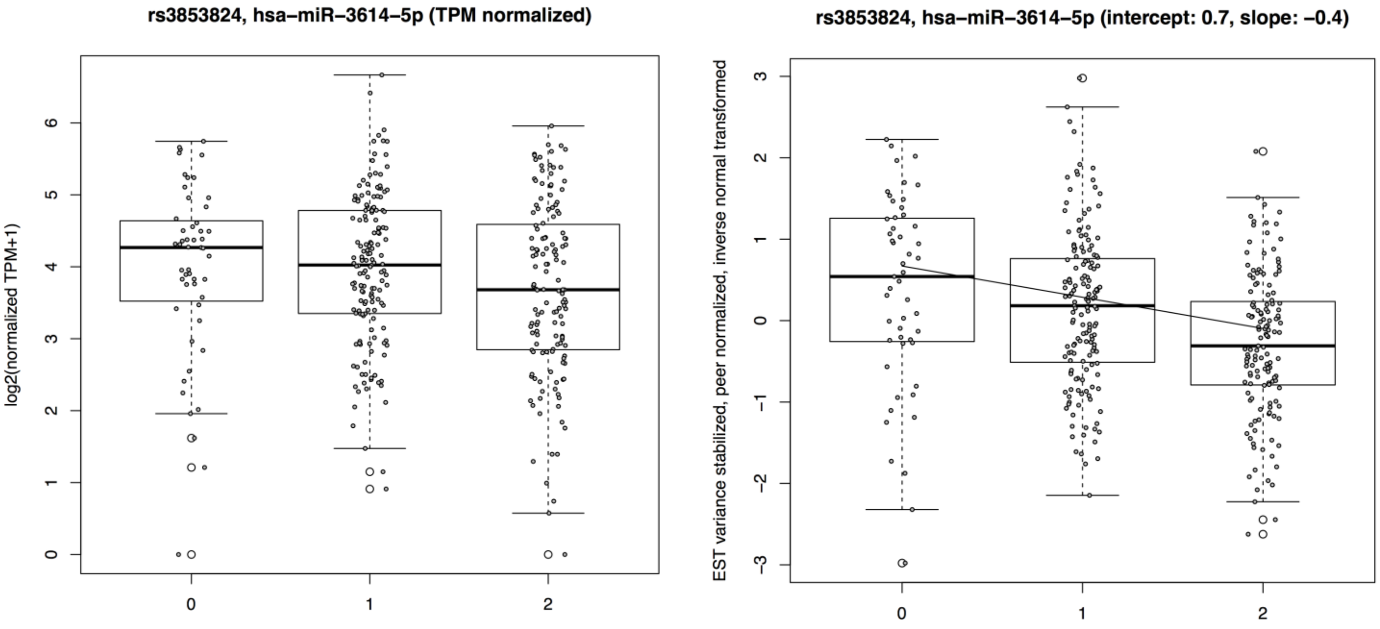

**Figure 9: hsa-miR-3614-5p gene expression levels according to genotypes of Crohn's disease risk SNP rs3853824.** See Suppl. Figure 2 for legend.

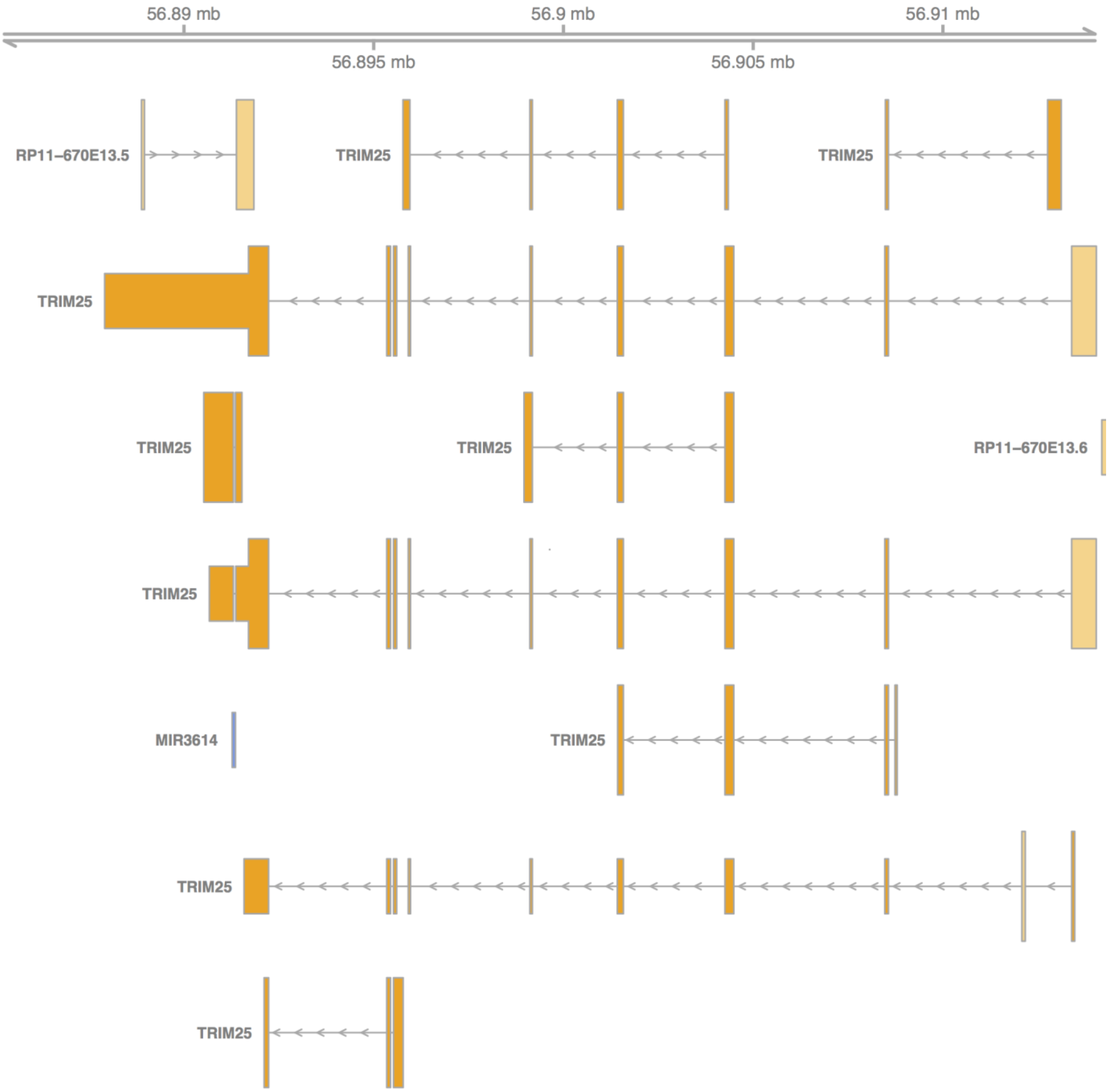

**Figure 10: Region of transcripts that overlap mir-3614.** Narrow boxes denote untranslated regions.

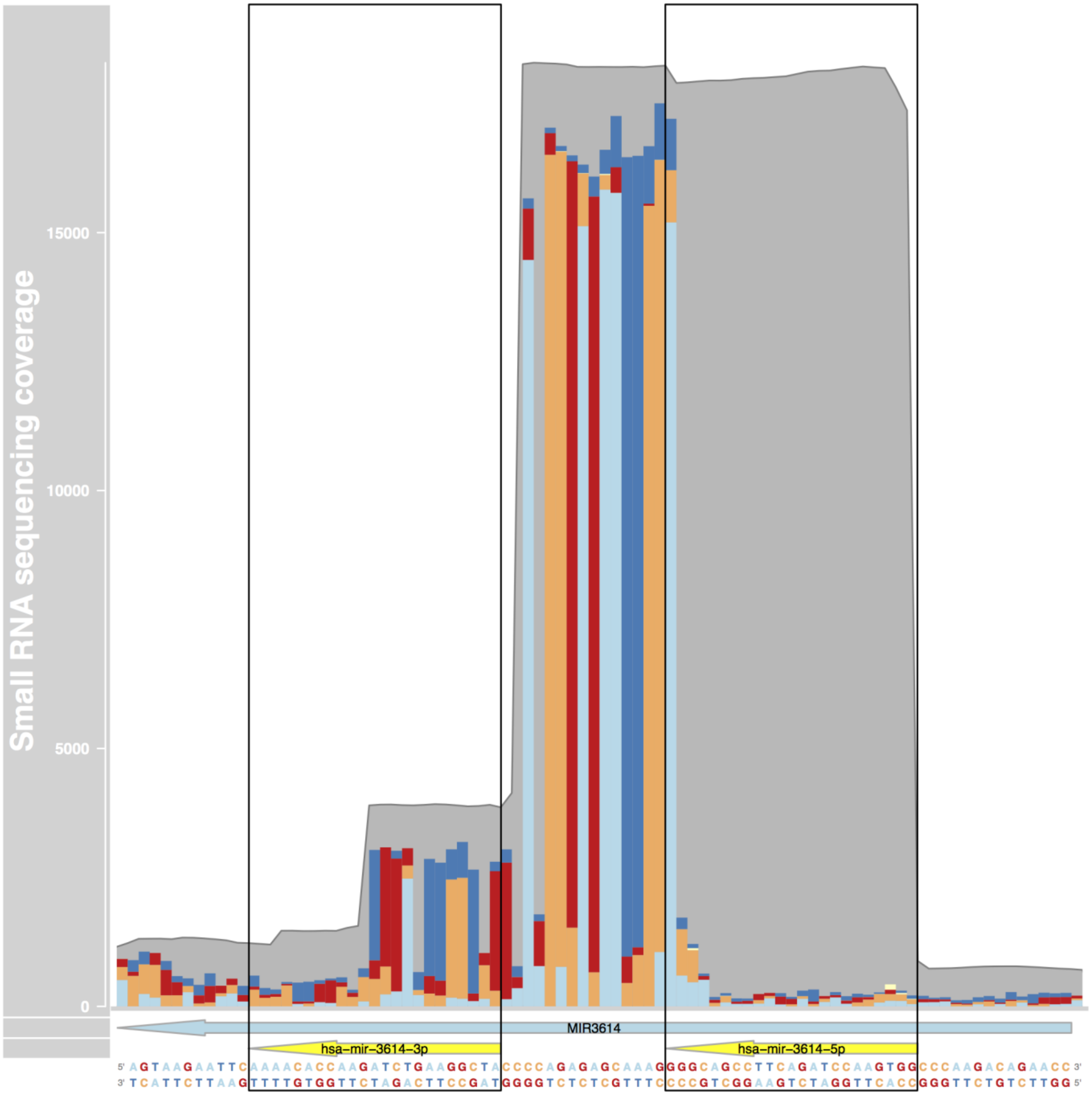

**Figure 11: Small RNA sequencing reads mapped to mir-3614 including the mature hsa-miR-3614-5p (right) and hsa-miR-3614-3p (left) miRNAs and to its surrounding sequence.** The mismatches downstream of the mature miRNA sequences can be attributed to Illumina primes and adapters, which also have been sequenced since read length is 36, which exceeds miRNA length.

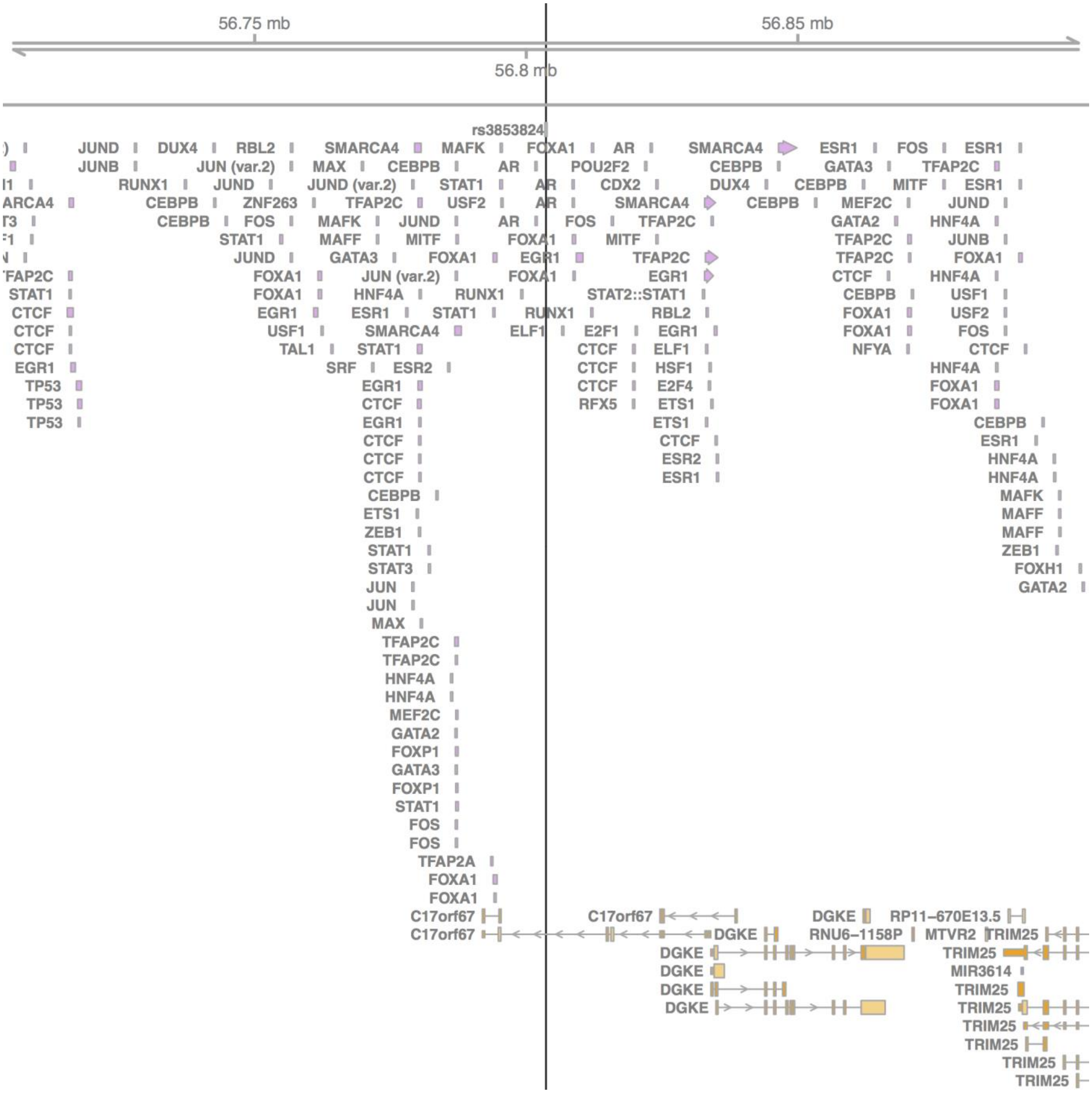

**Figure 12: SNP rs3853824 together with genes in the region (yellow) and regulatory regions from Oreganno labeled with corresponding targeting transcription factors (pink) and mir-3614.**

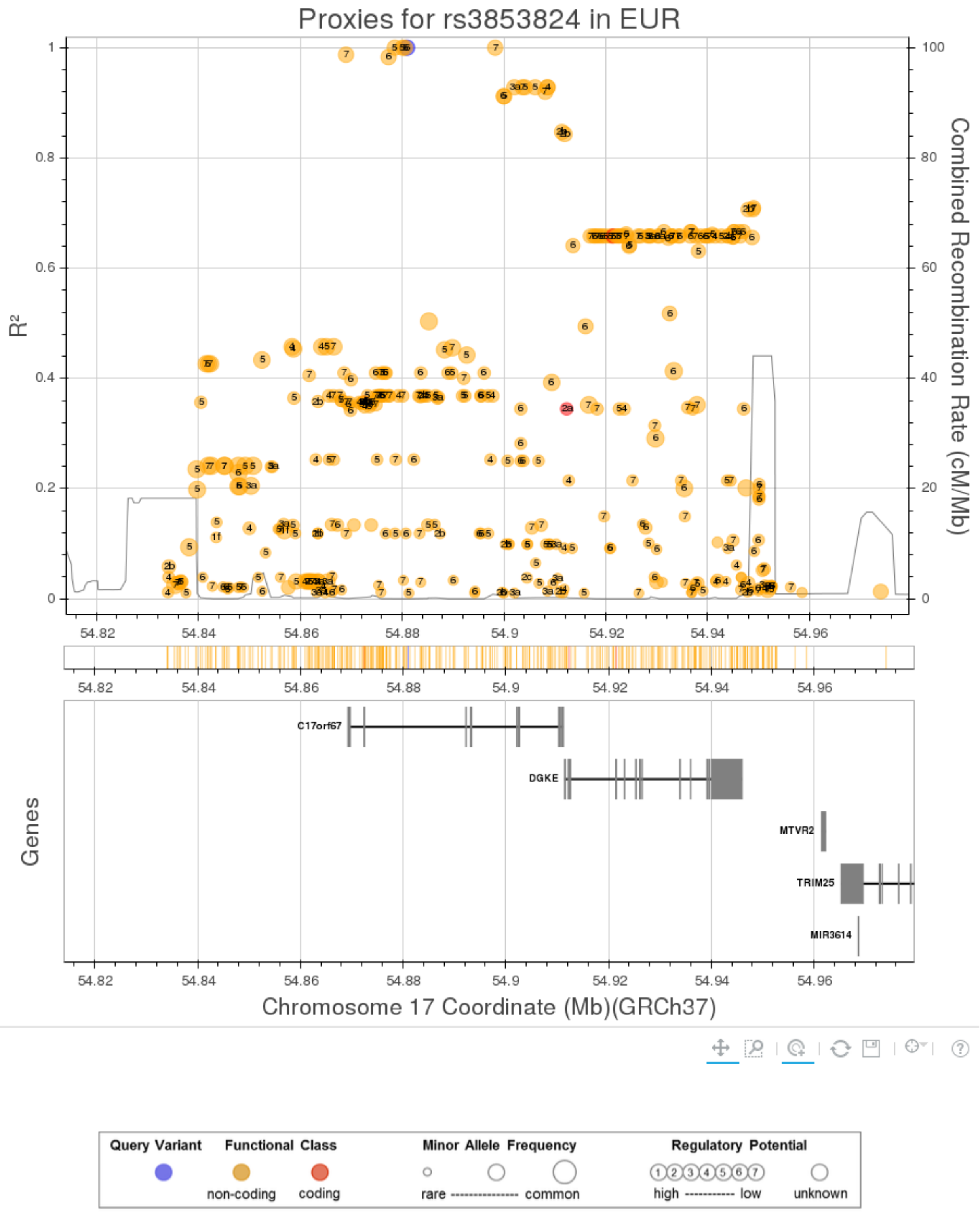

**Figure 13: Locus zoom plot for Crohn's disease index SNP rs3853824.** This figure has been generated using LDLink (https://analysistools.nci.nih.gov/LDlink) (19).

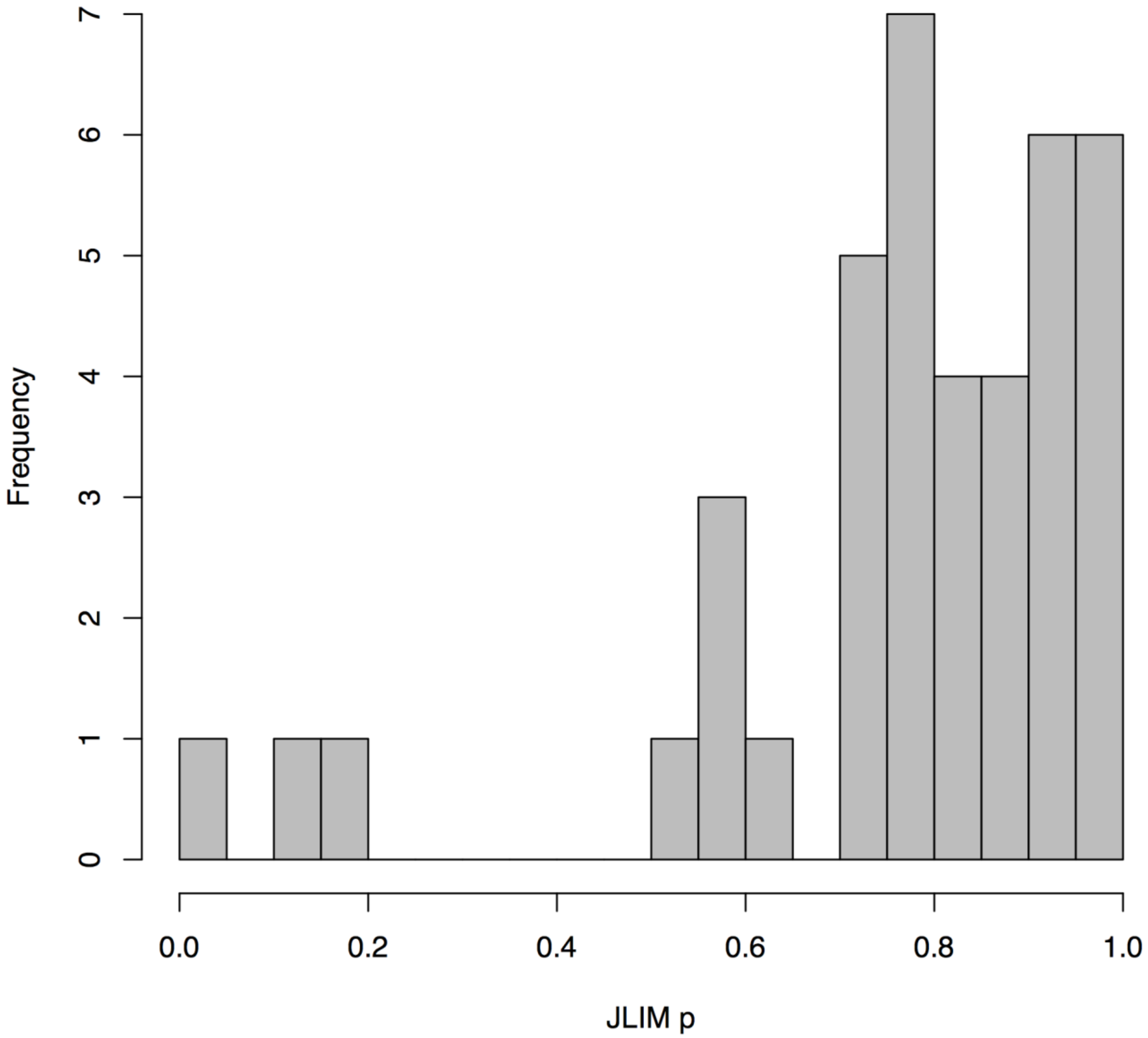

**Figure 14: Histogram of JLIM p-values.**
