## Supplementary figures and images for "Evidence for a potential role of miR-1908-5p and miR-3614-5p in autoimmune disease risk using genome-wide analyses" (from another version of this preprint: version 1, 2018-03-22)

### Supplementary Materials

#### Autoimmune disease-associated miRNA eQTL variants

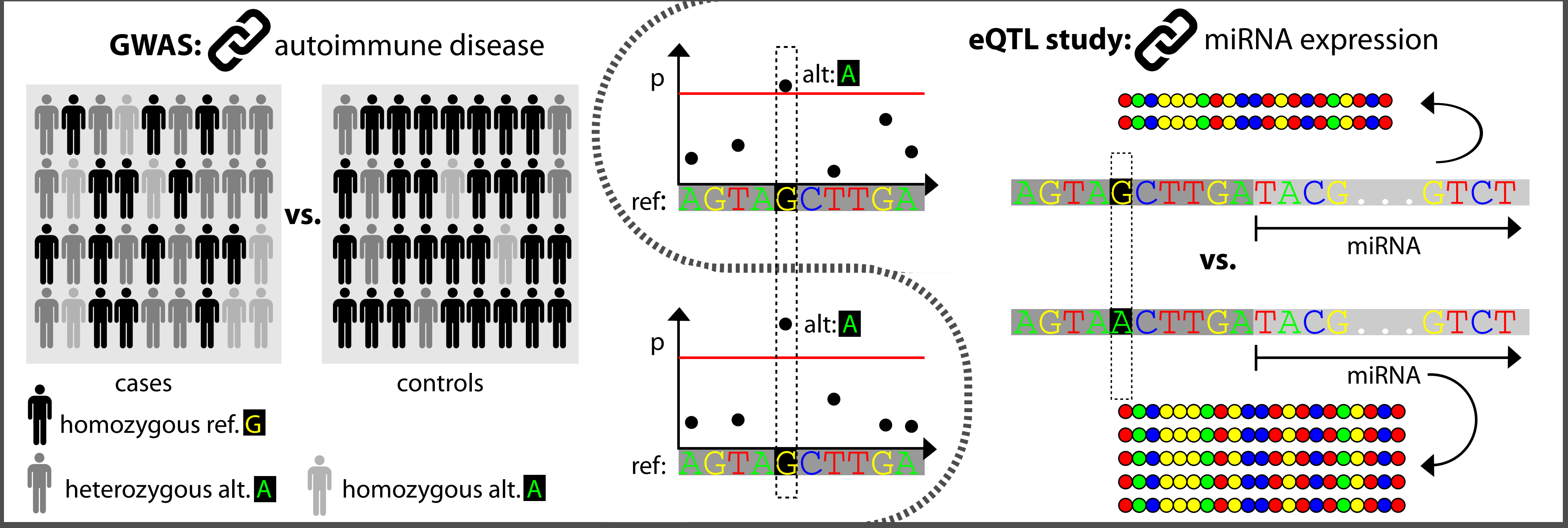
